# Quantitative analysis of nuclear pore complex organization in *Schizosaccharomyces pombe*

**DOI:** 10.1101/2021.08.11.455278

**Authors:** Joseph M. Varberg, Jay R. Unruh, Andrew J. Bestul, Azqa A. Khan, Sue L. Jaspersen

**Affiliations:** Stowers Institute for Medical Research, Kansas City, MO, United States; Department of Molecular and Integrative Physiology, University of Kansas Medical Center, Kansas City, KS, United States

**Keywords:** nuclear pore complex, nuclear envelope, Lem2, nucleolus, spindle pole body

## Abstract

The number, distribution and composition of nuclear pore complexes (NPCs) in the nuclear envelope (NE) varies between cell types and changes during cellular differentiation and in disease. To understand how NPC density and organization is controlled, we analyzed NPC number and distribution in the fission yeast *Schizosaccharomyces pombe* using structured illumination microscopy. The small size of yeast nuclei, genetic features of fungi and our robust image analysis pipeline allowed us to study NPCs in intact nuclei under multiple conditions. Our data revealed that NPC density is maintained across a wide range of nuclear sizes. Regions of reduced NPC density are observed over the nucleolus and surrounding the spindle pole body (SPB). Lem2-mediated tethering of the centromeres to the SPB is required to maintain NPC exclusion, which is important for timely mitotic progression. These findings provide a quantitative understanding of NPC number and distribution in *S. pombe* and show that interactions between the centromere and the NE influences local NPC distribution.

## Introduction

Nuclear pore complexes (NPCs) facilitate nucleocytoplasmic transport, organize the genome, influence gene expression and facilitate DNA repair (1–3). Each NPC is composed of multiple copies of *≈* 30 individual nucleoporin (Nup) proteins, which are organized around a central channel in eightfold symmetry (4–7). The NPC is anchored in the nuclear envelope (NE) by transmembrane Nups and through interactions between specific Nups and lipids of the nuclear membrane. Decades of research in a variety of systems has identified conserved functions for Nups in NPC assembly and transport and has mapped their organization within the structure of the NPC at nearly atomic resolution (8–11).

In contrast to our understanding of NPC structure, the mechanisms that control NPC density, distribution and composition remain poorly understood. Early studies using electron microscopy (EM) showed that NPC density is highly variable between species and cell types (12–14). NPC density does not appear to correlate with nuclear size or DNA content, but it is associated with metabolic activity (15) perhaps explaining links between changes in NPC density and cancer (16, 17) or in response to external signals (18–22). The remarkably long half-lives of many Nups (23–25) has led to a proposal that the number of NPCs in a cell is likely regulated at the stage of NPC assembly (reviewed in (26)). In metazoans and in budding yeast, the number of NPCs in the NE roughly doubles during interphase nuclear growth (21, 27–30). NPC assembly during the cell cycle is positively regulated by cyclin-dependent kinases (Cdks) (31) and negatively regulated by phosphorylation of NPC assembly factors by extracellular-regulated stress kinase (ERK) (32). However, some debate remains over the ubiquitous role of the ERK-mediated pathway for regulation of NPC density (33, 34).

Once assembled into the NE, NPCs adopt a variety of non-random distributions, ranging from pairs and clusters to higher-order linear and hexagonal arrays (12). Plant, animal and fungal nuclei have reduced NPC density in regions over the nucleolus and near sites of contact between the nucleus and cytosolic organelles, such as the vacuole/lysosome, Golgi apparatus and mitochondrion (12, 27, 35–40). Despite decades of work clearly demonstrating non-random NPC distributions in multiple cell types, little is known about how these patterns are formed and maintained. In metazoans, NPC distribution is mediated at least in part through the nuclear lamina (33, 41–43). However, as both plants and fungi lack lamins, additional factors must serve to regulate NPC distribution. LAP2-emerin-MAN1 (LEM)-domain proteins, which associate with the inner nuclear membrane (INM) throughout eukaryotes, are leading candidates; for example, emerin is enriched at pore-free regions of the NE in cultured cells (44). In budding yeast, NPC density is increased in the region of the NE near the spindle pole body (SPB), suggesting that either the SPB itself or associated factors control NPC recruitment or assembly (27, 39, 45).

Analysis of NPC composition in the region over the nucleolus in *Saccharomyces cerevisiae* showed that nucleolar-associated NPCs lack two Nups, Mlp1 and Mlp2 (46, 47). These orthologs of vertebrate Nup Tpr (translocated promoter region) are core structural components of the nuclear basket, a nucleoplasmic extension of the NPC that serves as a binding site for chromatin, proteasomes and other factors (48–51). Despite the clear evidence that *S. cerevisiae* maintain compositionally distinct populations of NPCs in subregions of the NE, we currently lack any mechanistic insight into how this is achieved. Further, it is unknown whether this is a unique property of budding yeast nuclei, as may be the case for other aspects of fungal NPC biology including mechanisms controlling inheritance of NPCs during mitotic divisions (which rely on the *S. cerevisiae* bud-neck structure) (52–56) and NPC remodeling during budding yeast meiosis (which is not seen in *S. pombe*) (57, 58). Identifying the mechanisms that control heterogeneity in NPC composition and distribution are of great interest, as transcriptomic and proteomic studies in metazoans have identified cell-type specific Nup expression patterns and shown that changes in NPC composition that are critically important for cell development, differentiation and progression of various diseases (59–64). These findings, in combination with the evidence for NPC compositional heterogeneity within individual nuclei in yeast, highlight the emerging concept that subpopulations of NPCs with distinct compositions and potentially specialized functions may exist at specific locations within the nuclear envelope (65).

Using *S. pombe* as a model system, we combined multiple quantitative imaging approaches, including threedimensional structured illumination microscopy (3D-SIM), to examine the number, distribution, and composition of NPCs in whole nuclei. We quantify NPC number under a range of conditions and show that fission yeast maintains a constant NPC density throughout its life cycle. NPC density appears to be maintained through a mechanism that links NPC assembly to available NE surface area. Experiments using 3D-SIM and live-cell imaging revealed a common structural organization of NPC clusters and identified two distinct behaviors of clusters during mitotic cell division. We show that the previously reported reduction of NPC density and alteration of NPC basket composition over the nucleolus-facing region of the NE is conserved in fission yeast. Additionally, NPCs are excluded from the NE region surrounding the SPBs by Lem2 and other factors.

## Results

### 3D-SIM image analysis pipeline for NPC quantitation

We developed an imaging and analysis pipeline to visualize and count NPCs in fission yeast following three-dimensional structured illumination microscopy (3D-SIM) of entire nuclei containing endogenously tagged Nups. This approach provides a roughly two-fold increase in resolution as compared to conventional light microscopy, with lateral resolution that approaches the size of the yeast nuclear pore (*≈* 90-120 nm diameter) (11, 66). Individual foci corresponding to single NPCs (FWHM = 121.6-136.5 nm, 95% CI) as well as larger foci that likely represent clusters of NPCs that cannot be fully resolved by SIM were detected throughout the nucleus (**Fig. 1A**). NPCs could be visualized using various tagged Nups, including representatives from each NPC subcomplex (**Fig. 1B**). NPC number and position, nuclear size and cell cycle stage could be extracted from images using the strategy outlined in **Fig. 1C** and these values could be used to derive measurements of nuclear surface area and NPC density (see Materials and Methods).

**Fig. 1.**
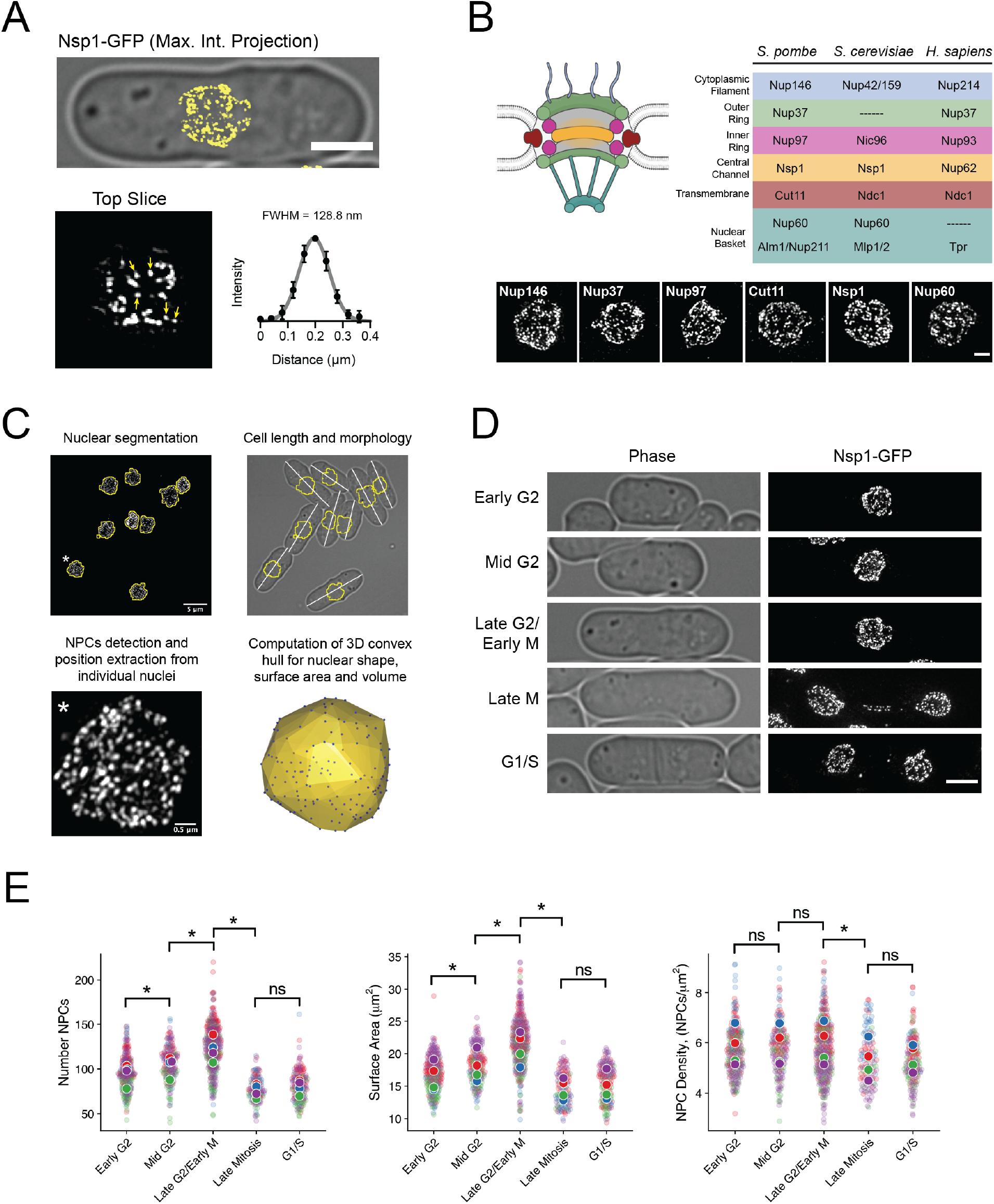
A 3D-SIM imaging and analysis pipeline to measure NPC number and density in *S. pombe*. A) 3D-SIM image of Nsp1-GFP overlayed on transmitted light image. Bar, 3 μm. Arrows show five foci representing single NPCs, which were fit to a Gaussian function to generate an average NPC intensity profile (FWHM, full-width half-maximum). B) NPC schematic with sub-complexes colored to match the table of individual Nups shown on right, with representative 3D-SIM images below. Bar, 1 μm. C) Pipeline for NPC analysis. D) Representative 3D-SIM image from each cell cycle stage. Bar, 3 μm. E) Mean number of NPCs, nuclear surface area and NPC density measurements from four independent replicates. Significant differences (*) determined using Wilcoxon rank sum tests. ns, not significant.

In *S. pombe*, nuclear size increases through interphase to maintain a constant nuclear-to-cell volume ratio (67). We observed that the number of NPCs also increases through interphase to maintain a constant NPC density (**Table S1, Fig. 1D-E**). We observed occasional differences in NPC density between mother and daughter nuclei produced by the symmetric mitotic division in *S. pombe*, reminiscent of the elevated NPC density observed for daughter nuclei produced by the asymmetric mitosis in *S. cerevisiae* (52). Visualization of Cdc7-GFP, a kinase that preferentially localizes to the “new” SPB during anaphase B (68), showed that the asymmetric NPC densities we occasionally observed is random with respect to the inheritance of the “old” or “new” SPB (**Fig. S1A**). During the late stages of *S. pombe* mitosis, a subset of NPCs localize to the membrane bridge where they facilitate active transport before being selectively disassembled to trigger localized NE breakdown and spindle disassembly (69–71). In agreement with these findings, we observed NPCs undergoing disassembly in the anaphase bridge region: NPCs contained transmembrane (Cut11) and structural nucleoporins (Nup37) but lacked the basket (Nup60) (**Fig. S1B**). Because of their dynamic nature, bridge NPCs were excluded from quantitative cell cycle measurements.

Despite the improved lateral resolution offered by SIM, clustering of NPCs and the comparatively reduced axial resolution likely leads to undercounting of NPCs using 3D-SIM. To determine the extent of undercounting, we applied our analysis pipeline to simulated datasets modeling a range of NPC densities (**Fig. S1C**). For simulated densities ranging from 2-5 μm^2^, the measured values for NPC density and NE surface area fell within 10% of the true simulated values; the percent error increased in a density-dependent manner, with values ranging from 10-30% for densities up to 10 NPCs/μ m^2^. Due to this undercounting, we used a secondary approach that did not rely on segmentation of individual NPCs from 3D-SIM images to measure NPC density. Comparison of nuclear size and total Nup-GFP intensities through the cell cycle showed similar increases in nuclear size and total NPC number, while NPC density (total Nup-GFP intensity per unit area) remained constant (**Fig. S1D**).

After correcting for the undercounting observed for our 3D-SIM approach, our analysis estimates that the average mid-G2 stage fission yeast nucleus contains between 115-137 ± 22-26 NPCs, with a nuclear surface area of 18.4 ± 2.8 μm^2^, for a density of 6.3-7.4 NPCs/μm^2^ (n=174). These values are lower than those reported for budding yeast (ranging from 9-15 NPCs/μm^2^, (13, 27, 72) but are similar to NPC densities reported for other cell types (ranging from 4.5-8 NPCs/μm^2^) (13, 14, 18, 30, 31). While similar trends in NPC number and density were observed using multiple tagged nucleoporins, variability in the number of NPCs detected was observed when comparing datasets between different tagged Nups (**Fig. S1E**). As a result, all experiments comparing genetic backgrounds or treatment conditions were performed in cells expressing the same tagged Nup.

### NPC density is controlled in a NE surface area-dependent manner

NPC density might be controlled by a mechanism coupling NPC assembly with available NE surface area (32). To explore this possibility in *S. pombe*, we examined NPC density in cells with nuclei covering a broad range of sizes. Meiotic progeny, known as spores, have a similar NPC density to mitotic cells despite having nuclei with 3-4 fold lower nuclear surface area (**Fig. 2A**) (Nsp1-mCh: Spores = 5.4 ± 1.2 NPCs/μm^2^, n=718; Mitotic, = 5.1 ± 0.93 NPCs/μm^2^, n=207). Similarly, a constant NPC density was maintained when nuclear size was reduced in mitotic cells using a temperature-sensitive allele of Wee1 kinase (*wee1*.*50*), a negative regulator of the cyclin-dependent kinase Cdk1/Cdc2 (73) (**Fig. 2B**) (Nup37-mCh: 25°C = 3.7 ± 0.86 NPCs/μm^2^, n=382; 36°C = 4.0 ± 0.95 NPCs/μ m^2^, n=506). Cells expressing a temperature-sensitive mutation in the Cdk1/Cdc2-phosphatase Cdc25 (*cdc25*.*22*) arrest at the G2/M boundary yet continue to increase both cell and nuclear size (74). During *cdc25*.*22* arrest, both nuclear surface area and the number of NPCs roughly doubled over the 3.5-hour incubation, allowing NPC density to be maintained (Nsp1-GFP: 25°C = 6.4 ± 1.3 NPCs/μm^2^, n=106; 36°C = 6.2 ± 1.0 NPCs/μ m^2^, n=63) (**Fig. 2C**). The increase in NPC number was dependent on NE membrane expansion during arrest, as chemical inhibition of fatty acid synthesis by treatment with cerulenin blocked nuclear growth while NPC density was maintained (36°C + Cerulenin = 6.8 ± 1.5 NPCs/μm^2^, n=110) (**Fig. 2C**). Yeast lacking core components of the autophagy machinery (*atg8*Δ or *atg1*Δ) (75) that targets NPCs for degradation during nutrient deprivation do not show increased NPC density compared to wild-type cells, suggesting that autophagy is not used to remove NPCs to maintain NPC density (**Fig. 2D**) (Nsp1-mCh: WT = 5.2 ± 0.83 NPCs/μm^2^, n=252; *atg1*Δ= 5.0 ± 0.87 NPCs/μm^2^, n=359; *atg8*Δ= 5.0 ± 0.97 NPCs/μm^2^, n=326). These results support a model whereby NPC density is maintained by a mechanism that restricts the assembly of new NPCs in the absence of increased available NE surface area.

**Fig. 2.**
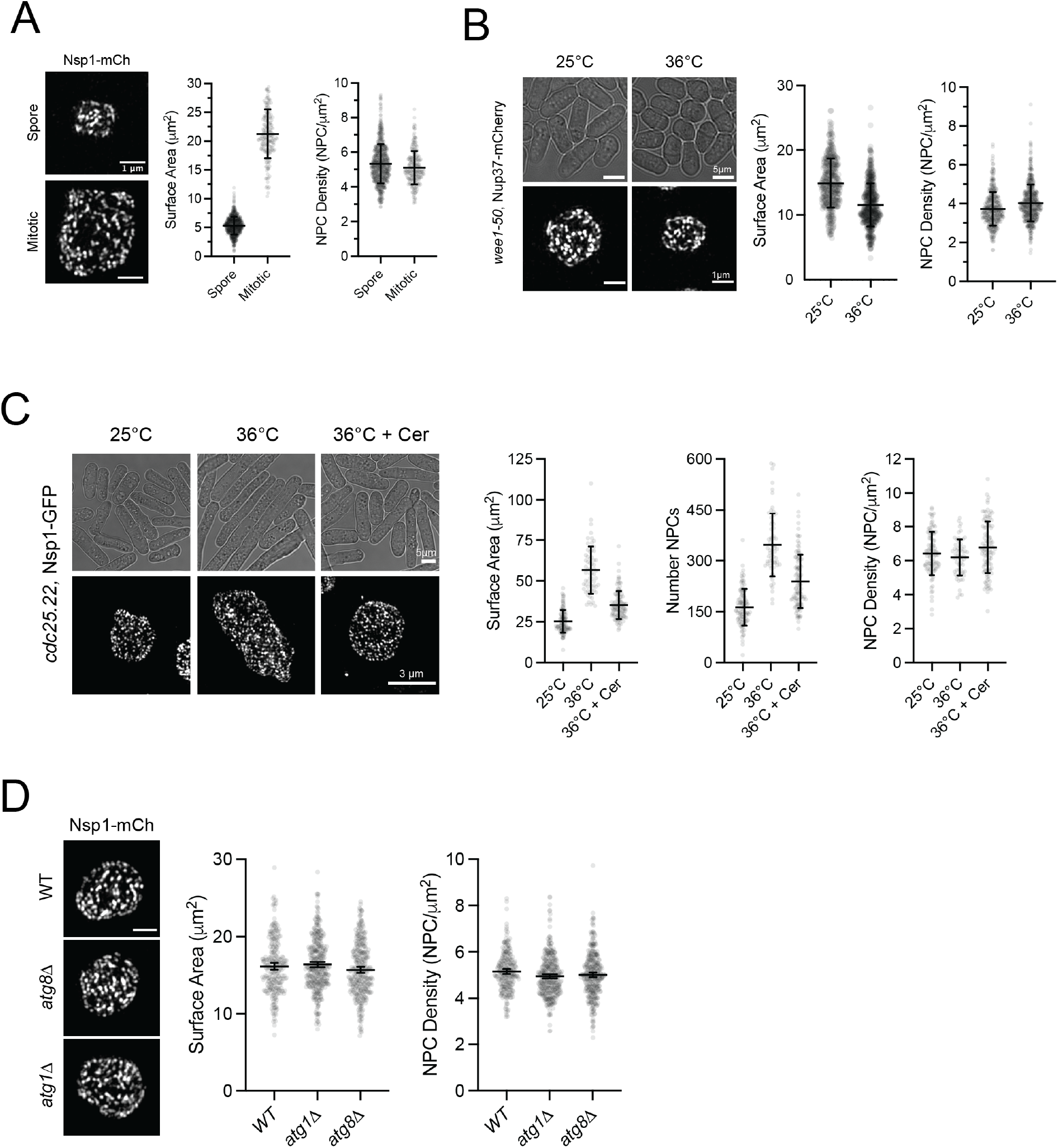
Surface area-dependent maintenance of NPC density. A) 3D-SIM image of Nsp1-mCh in nuclei from meiotic spores (top) or mitotic G2 (bottom) nuclei, with quantitation on right. B) 3D-SIM and quantitation of Nup37-mCh NPCs in *wee1*.*50* mutants grown at 25°C or shifted to 36°C for 3.5 h. C) 3D-SIM and quantitation of Nsp1-GFP in *cdc25*.*22* mutants at 25°C or shifted to 36°C for 3.5 h in the absence or presence of 10 μM Cerulenin (Cer). D) 3D-SIM and quantitation of Nsp1-mCh NPCs in wild-type, *atg8*Δ and *atg1*Δ cells. Bars, 1 μm.

### NPC cluster organization and dynamics

Our ability to observe NPCs throughout entire nuclei using 3D-SIM at near single-NPC resolution allowed us to evaluate higher level NPC organization. NPC clustering is common phenotype in different cell types and in mutants defective in NPC assembly. Using 3D-SIM, we compared NPC distribution in wild-type cells to two previously described clustering mutants: *nup132*Δ and *nem1*Δ (76–78).

Widefield and confocal images of NPC clusters in *nup132*Δ mutants often appear as one or two large clusters, however, 3D-SIM images revealed the presence of multiple smaller clusters distributed throughout the NE (**Fig. 3A**). The majority of *nup132*Δ nuclei displayed normally distributed NPCs or very mild NPC clustering, with only 14% displaying moderate to severe clustering (**Fig. 3A**). Unexpectedly, we frequently observed NPC clusters organized in a ring-like structure with diameters ranging from 250-300 nm (**Fig. 3B**). In rare cases, ring-like NPC clusters were also observed in wild-type cells, suggesting that these are not simply a unique phenotype of *nup132+* deletion. Clustering increased in aged *nup132*Δ cells grown on plates (**Fig. 3C**), consistent with previous reports (76). Similar rings were also observed in *nem1*Δ cells, which have increased rates of lipid synthesis that alters NE morphology and NPC distribution (70, 78–80) (**Fig. 3D**).

**Fig. 3.**
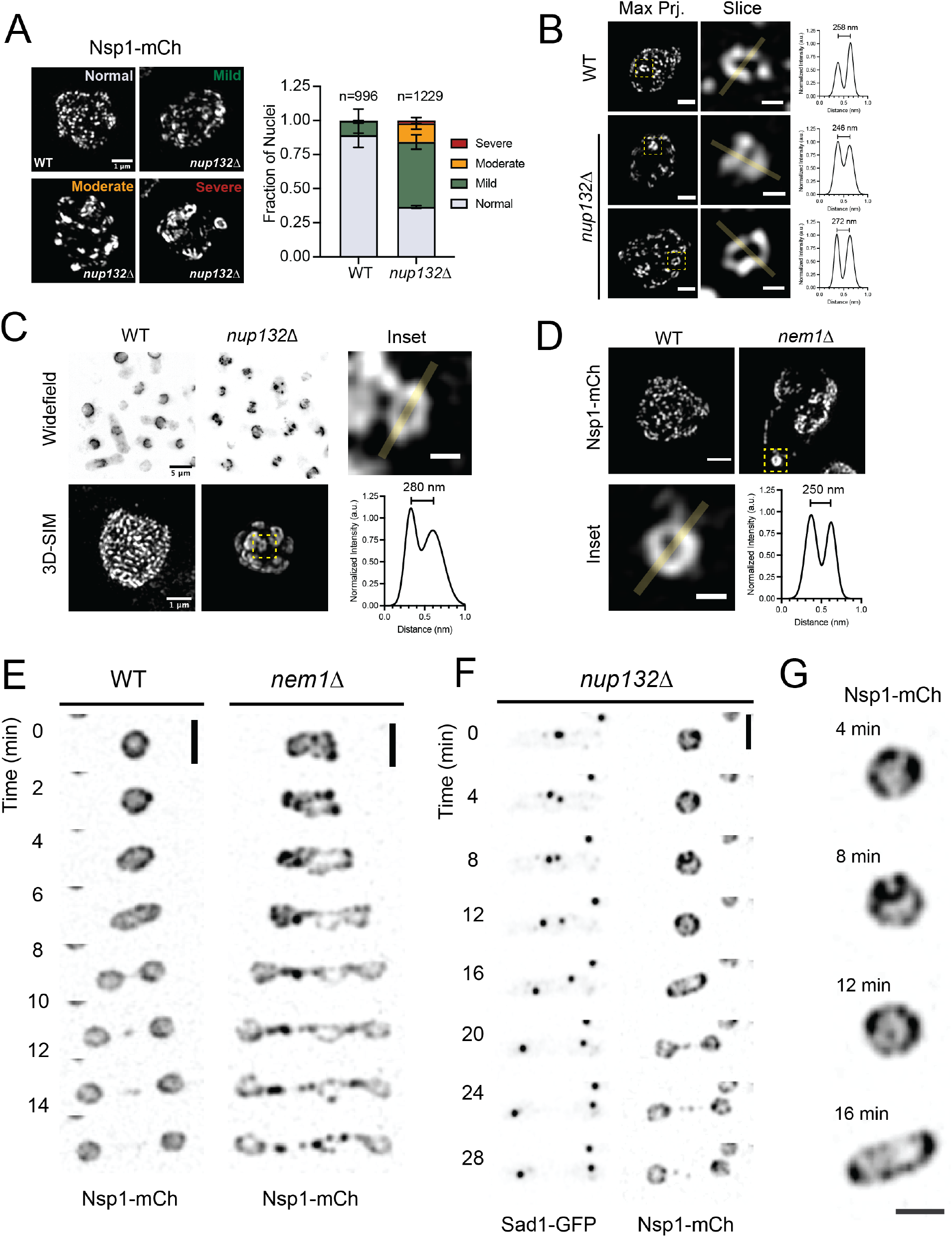
NPC cluster organization and dynamics. A) 3D-SIM of Nsp1-mCh in wild-type cells and in *nup132*Δ mutants. Quantitation of the frequency of clustering phenotypes from two independent replicates. Bar, 1 μm. B) Ring-like NPC clusters observed by 3D-SIM from cells in (A), including projections of the entire nucleus (left; bar, 1 μm) and magnified region (center; bar, 300 nm). Line profiles were taken across the indicated regions of the magnified region, with the corresponding intensity profiles shown at right. C) Clustering increases in *nup132*Δ cells grown on YES agar plates at 25°C for 7 d. D) 3D-SIM of Nsp1-mCh NPCs in wild-type and *nem1*Δ mid-G2 stage nuclei (Bar, 1 μm), with ring cluster shown in inset (Bar, 300 nm) and plot profile. E) Montage of time-lapse images of Nsp1-mCh in wild-type and *nem1*Δ cells. Bar, 5 μm. F) Montage of Nsp1-mCh and the SPB component, Sad1-GFP, in *nup132*Δ mutants. Nsp1-mCh intensity at the SPB relative to the average NE intensity is plotted, with the mitotic timepoints shown in the montage highlighted in gray. Bar, 5 μm. G) Insets of nuclei at the indicated time points from montage in F. Bar, 3 μm.

To examine the dynamics of the NPC clusters through the cell cycle, we performed time-lapse imaging of *nup132*Δ and *nem1*Δ cells and monitored NPC cluster dynamics in single cells. These experiments revealed two surprisingly different behaviors for clustered NPCs. In *nem1*Δ mutants, NPC clustering became more severe as nuclei prepared to divide. NPC clusters were frequently enriched in the anaphase bridge, along with excess membrane (**Fig. 3E, S2**). Following completion of nuclear division, the resulting daughter nuclei had normal NE morphologies and NPC densities equivalent to wild-type nuclei (**Fig. S2**). This suggests that *nem1*Δ nuclei can remove excess NE membrane and NPCs during mitosis via the anaphase bridge. In contrast, NPC clusters in *nup132*Δ nuclei coalesced into larger clusters that preferentially localized to the SPBs in mitosis (**Fig. 3G**). SPB-associated clusters are then segregated into the mother and daughter nuclei as cells complete mitosis. These observations show that at least two independent mechanisms exist to control NPC cluster dynamics and transmission during *S. pombe* nuclear division.

### Reduced NPC density and altered basket composition over the nucleolus

We observed a clear reduction in NPC density over the nucleolus (visualized using the RNA polymerase I subunit Nuc1) (81) from middle slices of 3D-SIM images (**Fig. 4A**), suggesting that a similar reduction of NPC density over the nucleolus occurs in *S. pombe* like *S. cerevisiae* (39). To quantitatively assess NPC density over this region we compared the intensities for multiple Nups from all subcomplexes over the nucleolus with intensities over the rest of the nuclear envelope (**Fig. 4B-C**). This confirmed that NPC density is reduced by *≈* 20% over the nucleolus-facing region of the NE and showed *≈* a 50% average reduction in the *S. pombe* Tpr orthologs Alm1 and Nup211 (48, 51, 82) (**Fig. 4D**). Like budding yeast, the NE region over the nucleolus has reduced NPC density and contains a unique population of NPCs that specifically lack the Tpr basket nucleoporins.

**Fig. 4.**
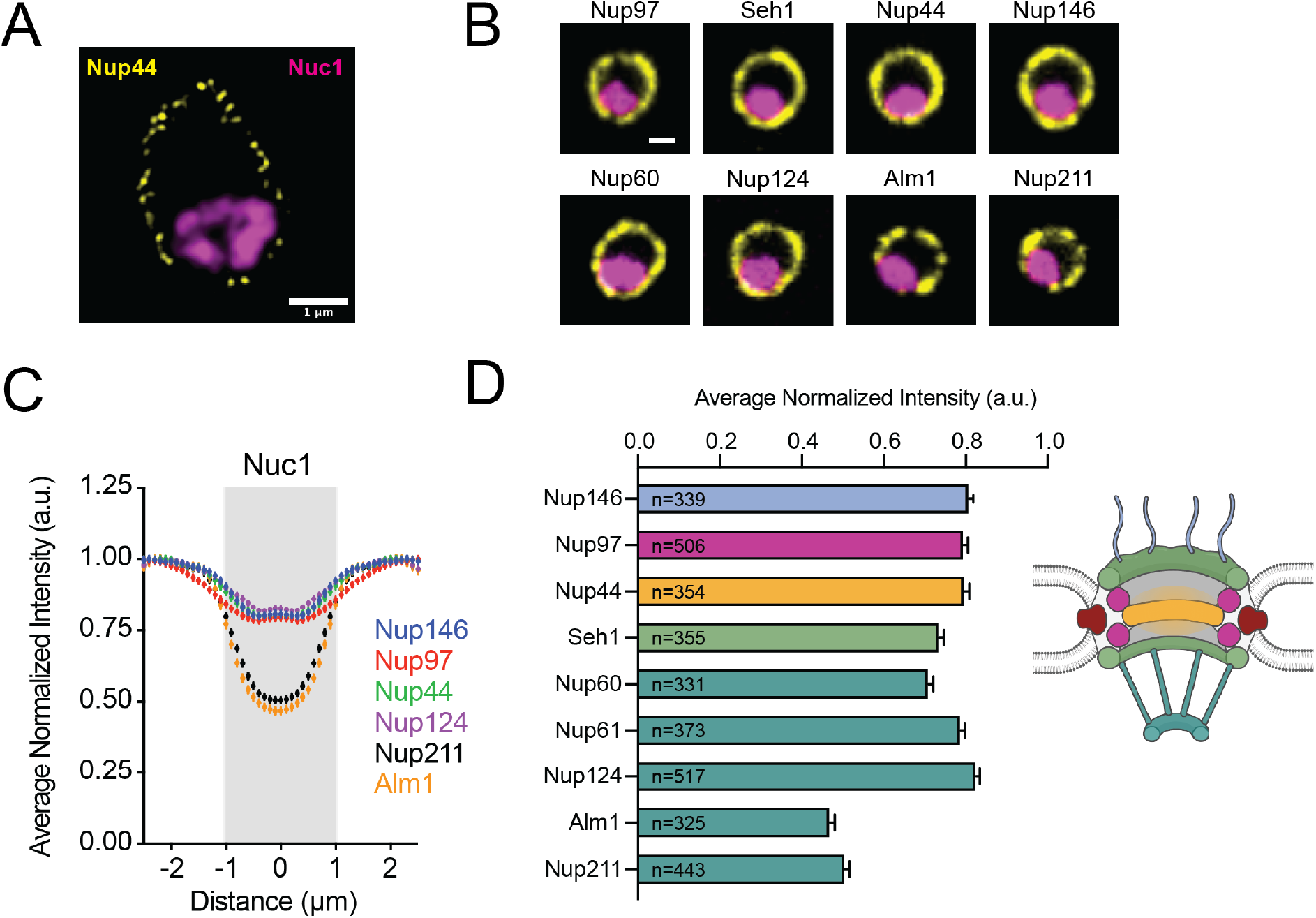
Reduced NPC density and altered NPC composition at the nucleolus. A) Middle slice from 3D-SIM image of Nup44-GFP NPCs and the nucleolus (Nuc1-mCh, processed with Gaussian blur). Bar, 1 μm. B) Confocal images of middle slices of nuclei showing Nups (yellow) and the nucleolus (magenta). Bar, 1 μm. C) Averaged profiles of Nup intensities at the NE relative to the nucleolus (gray, based on FWHM ± 95% CI of Nuc1-mCh). D) Average Nup intensity at position 0 in intensity profiles from (C), organized according to NPC subcomplex based on the schematic in Figure 1B. Error bars, SD. N, number of nuclei analyzed for panels C and D.

### NPCs are excluded from the SPB-proximal region throughout the mitotic cell cycle

In contrast to the nucleolar region, increased NPC density is found near the budding yeast SPBs (27), possibly due to a role of NPCs in NE remodeling during SPB insertion into the NE (45). EM analysis of fission yeast SPBs failed to identify an increased presence of NPCs within *≈* 200 nm of the SPB regardless of cell cycle stage (**Fig. 5A**). To examine the distribution of NPCs relative to the SPB at high resolution, we used single particle averaging of multiple 3D-SIM images (SPA-SIM); this approach has allowed us to visualize SPB-proximal proteins in budding and fission yeast (83–86). In SPA-SIM, the position of the two duplicated but unseparated SPBs are used as fiduciary points to realign images from multiple nuclei. To ensure that NPC distribution is visualized from a top-down perspective, we restricted our SPA-SIM analysis to SPBs that were centrally localized in the x-y plane with respect to the nucleus (see Materials and Methods). A composite image was then generated representing the average distribution of proteins of interest with respect to the SPBs.

**Fig. 5.**
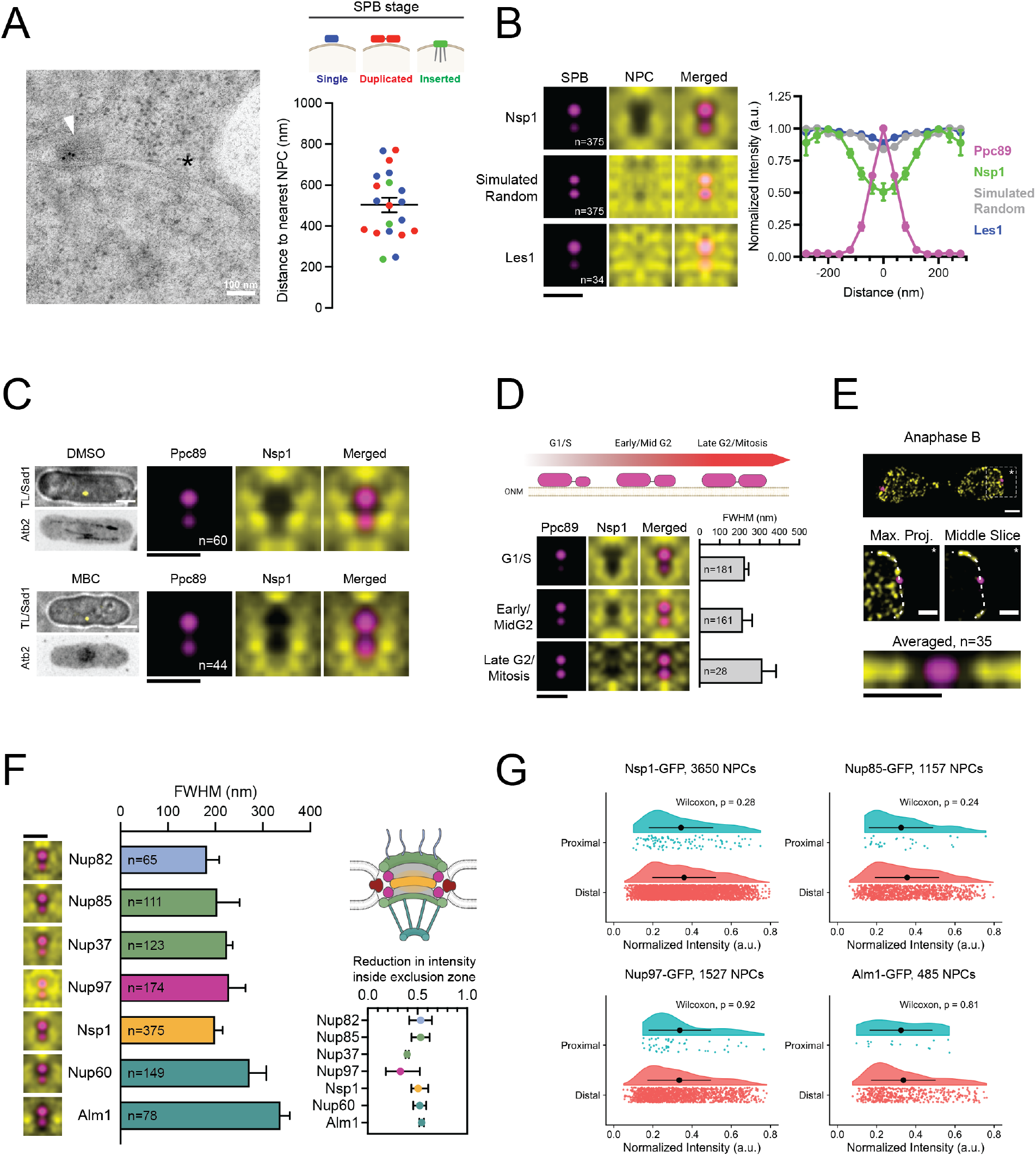
NPC exclusion from the SPB proximal region of the NE. A) ImmunoEM image of SPB component Ppc1-GFP (arrowhead). The nearest NPC is highlighted with asterisk. Plot of distance from SPB to the nearest NPC, based on SPB stage: blue = single SPB; red = duplicated SPB; green = inserted SPB. B) SPA-SIM images of Ppc89-mCh (magenta) and Nsp1-GFP (NPCs), Les1-GFP and simulated random distributions (yellow). Normalized intensity profiles across the mother and daughter SPBs. Error bars, SD. Bar, 0.5 μm. C) Confocal image for microtubules (mCh-Atb2) and SPBs (Sad1-GFP) in cells treated with DMSO (control) or 25 μg/mL MBC for 1 h to depolymerize microtubules. Bar, 3 μm. SPA-SIM images of Ppc89-mCh (magenta) and Nsp1-GFP (yellow) in these cells. Bar, 0.5 μm. D) Schematic of SPB duplication through the cell cycle. SPA-SIM images of Ppc89-mCh (magenta) and Nsp1-GFP (yellow) based on daughter/mother Ppc89-mCh intensity ratios (G1/S, 0.5; Early/Mid G2, 0.5 – 0.8; Late G2/Mitosis, 0.8). Plot of FWHM of Nsp1 exclusion zone for each stage. Error bars, 95% CI. Bar, 0.5 μm. E) 3D-SIM projection of Nsp1-GFP (yellow) and Ppc89-mCh (magenta) in anaphase. Bar, 1 μm. Enlarged images of SPB region, showing maximum projection and single middle z-slice. Bar, 0.5 μm. Averaged image of Nsp1-GFP NPCs relative to SPB in mitotically dividing nuclei (see Methods). Bar, 0.5 μm. F) SPA-SIM images of GFP-tagged Nups (yellow) and Ppc89-mCh SPBs (magenta), along with FWHM plot as in D. The majority of Nups have *≈*50% reduction in intensity near the SPB relative to the surrounding NE. Bar, 0.5 μm. G) Individual data points and kernel-smoothed density distributions of Nup-GFP intensities for NPC foci that were proximal (< 100 nm) or distal (>100 nm) to the SPB. Nup intensities for proximal and distal NPCs were compared using a non-parametric, unpaired Wilcoxon rank sum test. Black dots represent the mean normalized intensity value, and error bars show SD.

Consistent with EM analysis of NPC distribution around the fission yeast SPB, we observed a clear zone of NPC exclusion surrounding the SPBs in asynchronous populations of exponentially growing cells (**Fig. 5B**). This exclusion was not seen for Les1, an INM protein that is not a component of the NPC (70) or from simulations of randomly distributed NPCs (**Fig. 5B**). This exclusion zone was highly reproducible and cell cycle independent (**Fig. 5D-E**), with an average diameter of *≈* 200 nm (FWHM = 183.8-217.2 nm, 95% CI). We considered that exclusion of NPCs from the SPB could be due to forces exerted on the SPBs during interphase through the activity of microtubule-based motor proteins (87). How-ever, the exclusion zone was not altered following disruption of microtubules by treatment with the depolymerizing agent methyl benzimidazol-2-yl carbamate (MBC) (87, 88) (**Fig. 5C**).

We envisioned at least two possible models that could explain the NPC exclusion near the SPB. In the first, NPCs could be physically excluded from this region, perhaps through the presence of nuclear membrane proteins localized to the SPB region. This could include factors such as the SUN (Sad1-Unc-84 homology) domain-containing protein Sad1, which interacts with KASH (Klarsicht, ANC-1, Syne Homology) domain proteins Kms1 and Kms2 to form a LINC (Linker of Nucleoskeleton and Cytoskeleton) complex that tethers the cytosolic SPB to the NE (89, 90), or through proteins that tether centromeres to the NE (reviewed in (91)). Alternatively, the exclusion could represent a localized region of the NE containing NPCs with reduced Nup intensity, perhaps representing partial NPC disassembly or assembly intermediates.

Multiple lines of evidence suggest that the reduced Nup intensities near the SPBs is the result of physical exclusion of NPCs from this region. First, we observed similar exclusion patterns for multiple Nups including members of each NPC subcomplex (**Fig. 5F**). The size of the exclusion zone was relatively stable for all Nups, although was slightly larger for components of the nuclear basket (**Fig. 5F**). The majority of Nups were reduced by *≈* 50% over the SPB-proximal region, although the structural components Nup97 and Nup37 were excluded to a lesser extent (**Fig. 5F, inset**). If the observed Nup exclusion was due to changes in NPC composition near the SPB, we expected to see a relationship between NPC/Nup intensity and proximity to the SPBs. However, the intensities for individual NPCs that were proximal (within 100 nm radius of SPBs) and those that were distal were equivalent for multiple Nups, and only marginally (*≈* 20%) reduced for a subset of Nups (**Fig. 5G, Fig. S3**). Collectively, these findings support a model in which the reduced Nup intensity surrounding the SPBs is the result of reduced presence of NPCs in this region, rather than localized alterations of NPC composition.

### Exclusion of NPCs near the SPBs requires Lem2-mediated centromere tethering

We recently showed that the INM protein Lem2 localizes to the SPB during interphase and forms a ring with similar dimensions to that of the NPC exclusion zone (86, 92) (**Fig. 6A**). We hypothesized that the Lem2 ring may be a component of the physical barrier that prevents NPCs from localizing to this region. Indeed, deletion of *lem2+* resulted in a significant decrease in NPC exclusion from the SPB region (**Fig. 6B**) without affecting NPC composition, as SPB proximal and distal Nup intensities were similar in *lem2*Δ mutants. A decrease in NPC exclusion was not seen in cells lacking the INM protein Ima1 or the second *S. pombe* LEM domain-containing protein, Man1, that does not localize to the SPB (92) (**Fig. 6B**).

**Fig. 6.**
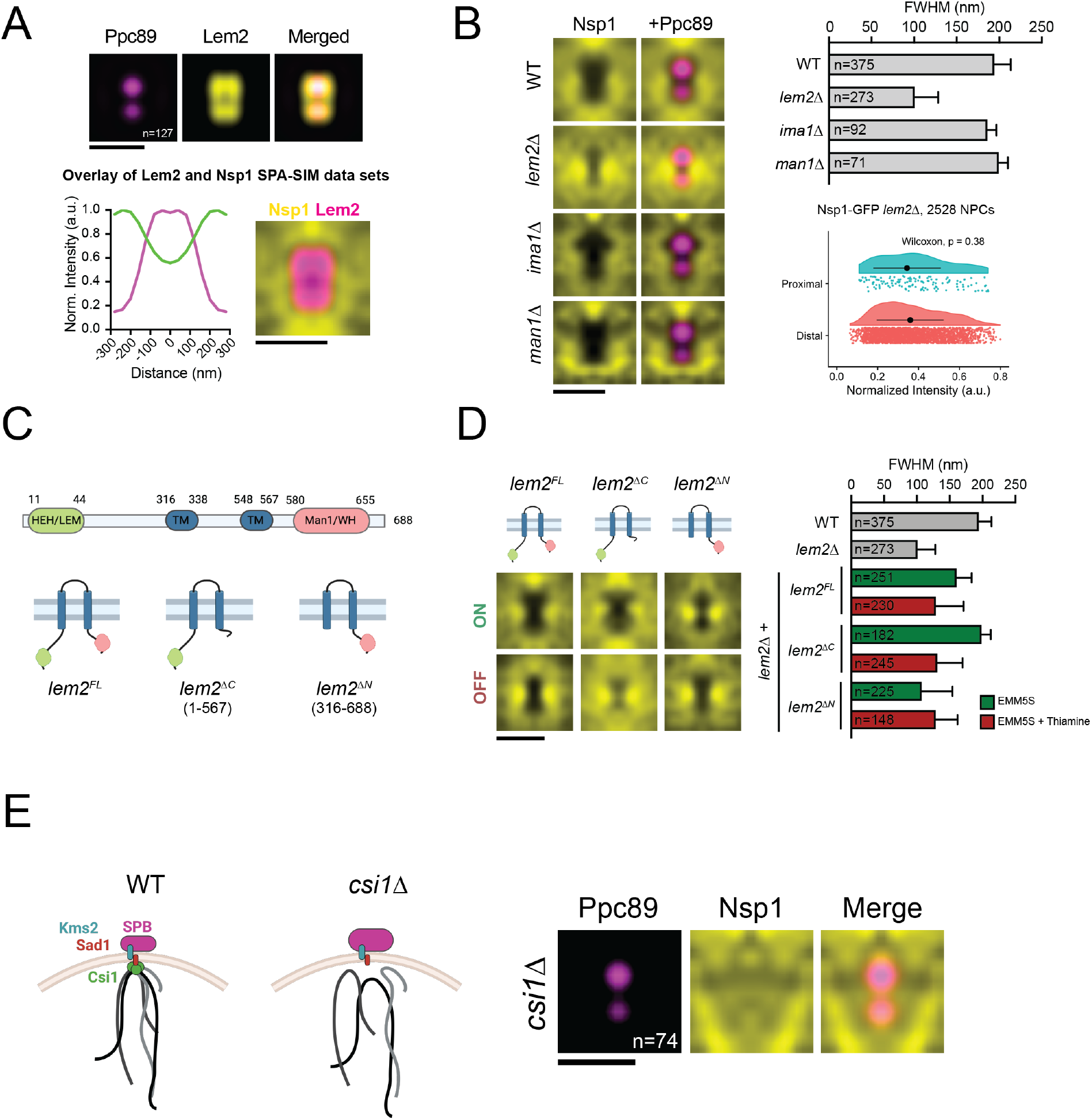
NPC exclusion requires Lem2 and centromere tethering. A) SPA-SIM of Lem2-GFP (yellow) and Ppc89-mCh (magenta). N, number of averaged images. Lem2-GFP and Ppc89-mCh intensity profiles are shown below. Overlay of SPA-SIM datasets for Lem2-GFP (magenta) and Nsp1-GFP (yellow). Bars, 0.5 μm. B) SPA-SIM Nsp1-GFP (yellow) and Ppc89-mCh (magenta) in wild-type, *lem2*Δ, *ima1*Δ and *man1*Δ backgrounds. Bar, 0.5 μm. Plot of NPC exclusion zone dimensions (FWHM, ± 95% CI). Nsp1-GFP intensity in SPB proximal and distal regions was plotted and compared using a Wilcoxon rank sum test. Black dots represent the mean normalized intensity value, and error bars show SD. C) Schematic of Lem2, including rescue constructs. D) SPA-SIM images of Nsp1-GFP (yellow) in *lem2*Δ cells with *lem2*^*FL*^, *lem2*^Δ*C*^ or *lem2*^Δ*N*^ constructs turned on or off. Bar, 0.5 μm. Plot of NPC exclusion zone dimensions (FWHM, ± 95% CI). For comparison, wild-type and *lem2*Δ dimensions from (B) are also shown. F) Schematic of centromere tethering at the SPB in wild-type and *csi1*Δ cells. SPA-SIM of Nsp1-GFP (yellow) and Ppc89-mCh (magenta) in *csi1*Δ cells. N, number of averaged images. Bar, 0.5 μm.

Lem2 contains two nucleoplasmic regions: an N-terminal HEH/LEM domain that is required for DNA binding and centromere tethering at the SPB (93, 94), and a C-terminal Man1/Winged-Helix domain that tethers telomeres to the nuclear periphery (95) (**Fig. 6C**). Lem2 truncation mutants lacking the N- and C-termini localize to the SPB (94), allowing us to test which regions of Lem2 are needed for NPC exclusion. Full-length or mutant versions of Lem2 were stably integrated at the *ura4+* locus and expressed as C-terminal 3xHA fusion proteins in a *lem2*Δ background using the thiamine-regulatable *nmt41* promoter system (96, 97) (**Fig. 6D, Fig. S4**). Exclusion of NPCs from the SPB region was similar to wild-type cells when either full-length (*lem2*^*FL*^) or *lem2*^Δ*C*^ constructs were expressed (EMM5S). However, *lem2*^Δ*N*^ expression resulted in an exclusion zone FWHM similar to *lem2*Δ mutants (**Fig. 6D**), suggesting that NPC exclusion depends on the function of Lem2’s DNA-binding N-terminal HEH/LEM domain.

The size of the NPC exclusion zone was reduced in *lem2*Δ and *lem2*^Δ*N*^ strains, however, NPCs were still strongly excluded from a smaller region directly underneath the SPBs. During interphase, fission yeast centromeres tether under the SPBs (98) through interactions with multiple proteins including Lem2, Sad1, and Csi1 (reviewed in (91). We hypothesized that the smaller exclusion zone observed in the absence of Lem2 could reflect a physical barrier formed by the remaining NE-centromere interactions. To test whether tethering of centromeres to the SPBs drives exclusion of NPCs from this smaller region, we examined NPC exclusion in *csi1*Δ mutants, in which *≈* 70% of cells exhibit defects in centromere tethering (99). Interestingly, in *csi1*Δ cells, NPCs were no longer excluded from the SPB-proximal region (**Fig. 6E**). This supports a model whereby the exclusion of NPCs from the SPB region is the result of physical interactions between centromeres and INM proteins, including Lem2, that tether the centromeres under the SPBs during interphase.

### Forcing NPCs into the SPB proximal region delays mitotic progression

A key question is why NPCs are excluded from the region near the SPB in *S. pombe*: is this important for SPB insertion into the NE during mitosis, for SPB separation or for some unknown aspect of SPB biology? Although *lem2*Δ and *csi1*Δ cells lose NPC exclusion at the SPB, these mutants likely have pleiotropic effects due to their multiple functions in nuclear size regulation, SPB function and chromosome organization (95, 99–104). Therefore, we developed an ectopic system to forcibly tether NPCs at the SPB (**Fig. 7A**). When expressed in mitotically growing cells, the INM protein Bqt1 localizes to the SPB through interactions with the INM protein Sad1 (93). Expression of Bqt1-mCh fused to GFP-binding protein (Bqt1-mCh-GBP) using a thiamine-repressible promoter (*nmt3X*) forced the recruitment of NPCs containing Nsp1-GFP (a component of both the central channel and the nuclear basket (105), into the SPB proximal zone, reducing the width of the exclusion zone by *≈* 50% (**Fig. 7B**). Forced localization of NPCs into the SPB proximal region caused a moderate impairment in growth (**Fig. 7C**). Using live cell imaging to visualize SPBs and microtubules in the presence and absence of Bqt1-mediated NPC tethering, we showed that this growth delay is likely the result of two defects. First, the time between the depolymerization of interphase microtubules and the formation of 3 μm metaphase spindle (106) was increased by *≈* 20% when NPCs were tethered (**Fig. 7D**). Second, NPC tethering caused spindle orientation defects, with prophase and metaphase spindles having an average deviation of *≈* 30 degrees from the longitudinal axis of the cell compared to *≈* 18 degrees when the Bqt1-GBP tether was repressed (**Fig. 7E**). Tethering did not affect microtubule nucleation at the SPB, including the formation of cytoplasmic microtubules. Together, these data suggest an unexpected role for a zone of NPC exclusion at the SPB that is important for mitotic progression.

**Fig. 7.**
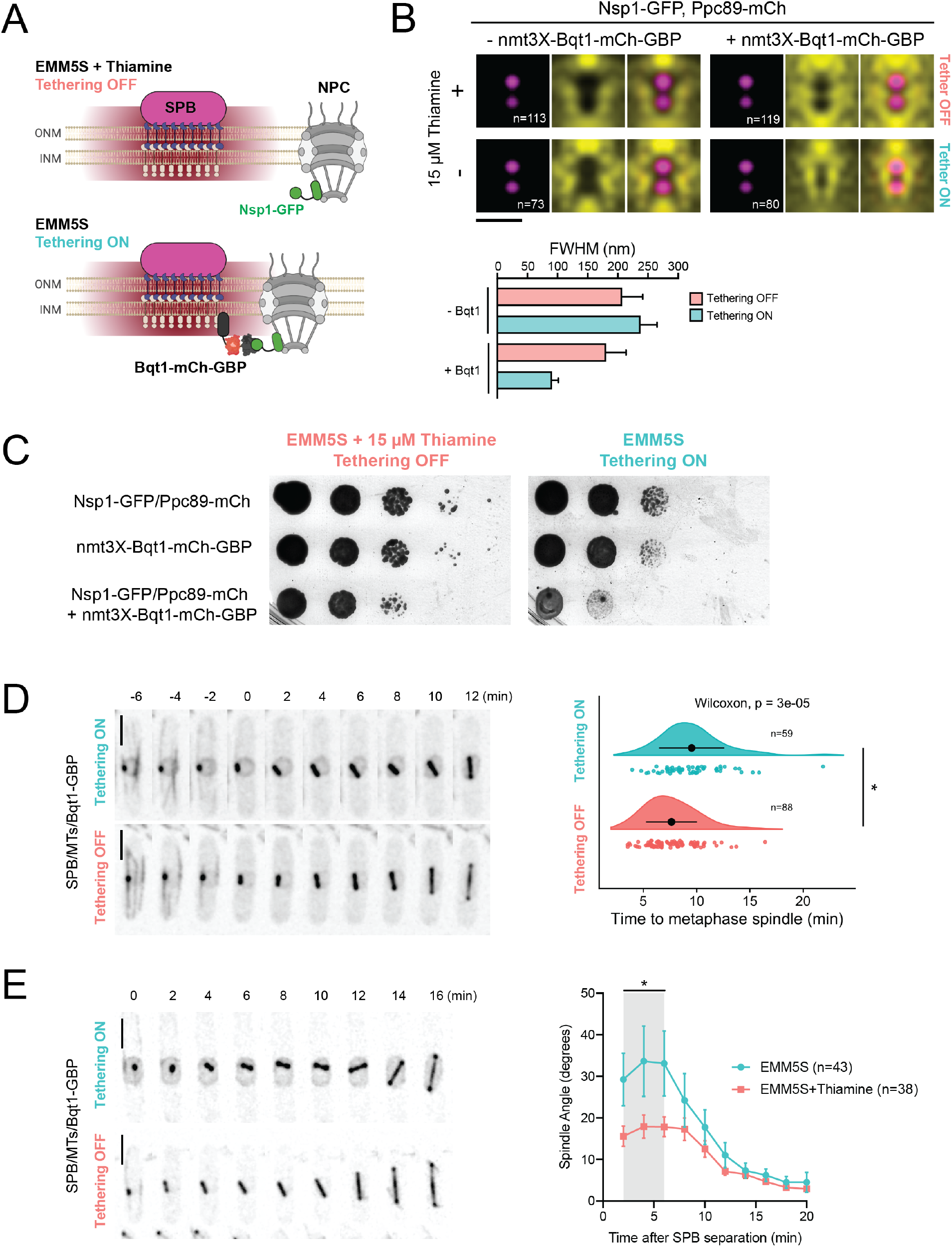
Loss of NPC exclusion at the SPB delays mitotic progression. A) Schematic of NPC tethering to the SPB. Bqt1-mCh-GBP localizes to the SPB and recruits NPCs via Nsp1-GFP into the SPB proximal region (red gradient). Bqt1-mCh-GBP expression is repressed by addition of thiamine to the media. B) Cells with or without Bqt1-mCh-GBP were grown in the presence (off) or absence (on) of thiamine. SPA-SIM images of Ppc89-mCh (magenta) and Nsp1-GFP (yellow). Bar, 0.5 μm. n, number of images. Dimensions of the NPC exclusion zone in each condition are shown below. Error bars represent 95% CI of SD of Gaussian fit to the Nsp1-GFP intensity profile. C) Growth of the indicated cells was tested at 30°C. D-E) Time-lapse imaging of Nsp1-GFP/Bqt1-mCh-GBP cells containing Ppc89-mCh (SPB) and mCh-Atb2 (microtubules). Time points are indicated above, with t=0 based on cytoplasmic microtubule disassembly. Bar, 5 μm. D) The time from cytoplasmic microtubule disassembly to formation of a metaphase spindle (3 μm) was analyzed and is shown on the right. N, number of cells analyzed. Statistical significance (*) was determined using a Wilcoxon rank sum test. Black dots represent the mean and error bars show SD. E) Plot of average spindle angle relative to the cell length axis versus time after SPB separation for the indicated number of cells (n). Error bars, SD. Timepoints highlighted in gray with asterisk determined to be significantly different between conditions using unpaired t-test with Welch correction.

## Discussion

Multiple imaging approaches, including EM and fluorescence microscopy, have been used to determine the number and distribution of NPCs in various systems. The higher resolution afforded by EM and super-resolution light microscopy methods often comes at a price of significant increases in the time required for sample preparation, image acquisition and analysis. In contrast, 3D-SIM generates high-resolution datasets using standard fluorescence microscopy approaches, allowing for quantitative analysis of NPC organization through whole nuclei. We apply 3D-SIM to fission yeast nuclei to provide the first map of NPCs in this system. We find that NPC density in *S. pombe* nuclei is similar to that described for many metazoan nuclei and is constant over a range of nuclear sizes. This finding has important implications regarding the mechanisms used for NPC assembly in fission yeast. For example, the total number of NPCs present in the two daughter nuclei in late mitosis is *≈* 26% greater than the number present in the mother nucleus prior to division. In agreement with previous findings (67), we observed that although the combined nuclear volume of the daughter nuclei is similar to the total volume of the mother nucleus, the combined surface area is *≈* 34% greater than that of the mother nucleus (**Table S1**). Together, this suggests that NPC assembly continues to occur during the rapid expansion of the NE during cell division (107). However, NE expansion during mitosis takes place over a time frame of roughly 20-25 min, significantly shorter than the 45-60 min required for completion of NPC assembly in budding yeast and during interphase in metazoans (29, 108). During the short cell cycles of the syncytial nuclear divisions in Drosophila embyros, rapid NPC assembly occurs via incorporation of assembly intermediates from annulate lamellae (109, 110). However, by EM and by fluorescence microscopy, we and others do not observe pools of NPCs/Nups outside the NE so it is unclear how *S. pombe* maintains NPC density during mitosis. Continued NPC assembly in *cdc25*.*22* arrested cells that have low Cdk1 activity suggests that unlike in metazoans, Cdk1 is not required for NPC assembly in *S. pombe*. The fact that the nuclear size increase and NPC assembly in *cdc25*.*22* were blocked by inhibiting fatty acid synthesis supports a model for negative regulation of NPC assembly similar to that proposed in vertebrates mediated by Tpr/ERK. In this model, signals emanating from existing NPCs inhibit assembly of NPCs in the surrounding region and inhibition of NPC assembly can be overcome by reducing NPC density through NE expansion. Spk1, the fission yeast ERK ortholog, is not essential for cell viability (111), suggesting that mechanistic details of this pathway may differ between species.

A major benefit of the 3D-SIM approach is that the improved resolution allowed for identification of distinct patterns of organization for clustered NPCs in *S. pombe*. NPC clusters were often observed to be organized in ring-like patterns *≈* 250-300 nm in diameter that were more prevalent in the clustering mutants *nup132*Δ and *nem1*Δ. These rings are smaller than typical yeast autophagosomes (300-900 nm diameter) (112, 113). although they are similar in size to nuclear-derived vesicles containing NPCs seen in EM images of NPCs being removed by autophagy in budding yeast (114). Our observation of increased ring number by 3D-SIM when *nup132*Δ cells were grown on solid instead of liquid media suggests that changes in nutrient availability triggers NPC reorganization into ring clusters of a consistent size. Whether the formation of these rings promotes their subsequent removal via autophagy or other pathways remains to be tested. However, the increased frequency of ring clusters in *nup132*Δ cells may provide insights into the mechanism driving their formation. Nup132 is a structural Nup that facilitates interactions between the structural scaffold of the NPC with lipids through its N-terminal ALPS motif (115, 116). Deletion of Nup132 may alter the interactions between NPCs and specific lipid species present in the NE, making *nup132*Δ cells especially sensitive to changes in lipid composition that may occur due to nutrient availability during growth on plates.

Time-lapse imaging of NPC clusters revealed two strikingly different approaches to how clustered NPCs are handled during mitosis. In *nem1*Δ mutants, both excess nuclear membrane material and NPCs are segregated into the anaphase bridge region during nuclear division (**Fig. 3F**), a distinct nuclear compartment that is a unique site of NPC disassembly (70, 71). Clusters of NPCs formed in *tts1*Δ cells specifically during mitotic NE expansion also localized to the anaphase bridge (117). This suggests that the anaphase bridge region of the NE may serve as a site where NPCs and NE material are sent to be removed during division, analogous to the NE-derived compartment that forms in budding yeast meiosis II to sequester and degrade NPCs (57, 118). The fact that *nup132*Δ clusters do not similarly localize to the anaphase bridge suggests that the fate of NPC clusters likely depends on the mechanisms driving the clustering. If *nup132*Δ clusters interact with specific lipids, this may also explain their portioning with the SPB, which has been proposed to contain a unique NE (119) (below).

Our results clearly demonstrate that the region of the NE over the nucleolus and near the SPB are distinct from other NE regions. It is not surprising that the nucleolar region has reduced NPC density and pores lacking the basket Nups Alm1 and Nup211 given similar observations of NPC number and composition in plants, mammals and fungi (12, 35, 36). In contrast, we were somewhat surprised to see reduced NPC density at the SPB in fission yeast given that NPC density is increased near SPBs in budding yeast (27, 45). Perhaps this is reflective of differences in the roles NPCs play in SPB insertion into the NE in the two fungi – NPCs are thought to facilitate SPB incorporation into the NE in *S. cerevisiae* but do not appear to be required for SPB assembly into the NE in *S. pombe*. Moreover, our data indicate that tethering of NPCs into the SPB proximal region in fission yeast delays the formation of a properly oriented metaphase spindle. This phenotype may be due in part to the artificial tether, although it is important to note that *csi1*Δ mutants, which lose NPC exclusion, also have mitotic defects (100, 120).

A key question is how populations of distinct NPC composition are established and maintained in specific regions of the NE, since fungal NPCs laterally diffuse through the NE. At least three potential models exist: intact NPCs diffuse into a subregion and are partially disassembled; a unique NPC subpopulation is assembled in that region of the NE; or sub-regions of the NE have unique properties and preferentially allow for NPCs of specific composition to diffuse in and/or be retained. Partial disassembly of fungal NPCs has been reported in multiple cell types and conditions (121–123). However, the fact that Nup exclusion is decreased at the SPB when NPCs are artificially tethered suggests that an SPB-derived signal such as phosphorylation by a SPB localized kinase likely does not induce NPC remodeling in this region of the NE. Instead, we favor a model for physical NPC exclusion from the SPB involving both centromeres and Lem2. At the SPB, tethering of centromeres to INM-localized SPB components forms a physical barrier that prevents the diffusion of NPCs through the NE into the SPB proximal region. NPCs that contain a basket may be particularly affected as they are likely to associate with chromatin and other complexes through this structure. Consistent with the steric model, if we reduced or eliminated centromere tethering, either by removing Lem2’s N-terminal HEH/LEM domain or by deletion of *csi1+*, the NPC exclusion zone was diminished. Interestingly, NPCs remain excluded from the SPB region throughout mitosis, including during periods where Lem2 no longer localizes to the SPB (92). During these stages, NPC exclusion is likely maintained by multiple proteins that form SPB-ring structures during mitosis, including Ima1 and Sad1 (86).

It is likely that the reduced NPC density and altered basket composition over the nucleolus is produced through a different mechanism. In budding yeast, the NE over the nucleolus is more amenable to membrane expansion than regions outside of the nucleolus (124). Similar differences in NE membrane properties over the nucleolus may exist in *S. pombe* and could drive the observed NPC heterogeneity. For example, differences in membrane composition or fluidity could alter the ability for NPCs to diffuse laterally through this portion of the NE, leading to reduced density over the nucleolus. Alternatively, the region over the nucleolus could have higher rates of NE membrane incorporation and NPC assembly. In this scenario, the reduced presence of Alm1 and Nup211 could be due to these Nups being the last components added during NPC assembly (108). In either case, the reduced presence of Alm1/Nup211 over the nucleolus could be the result of interactions between chromatin and NPCs containing Alm1/Nup211 (either directly or indirectly via basket-associated complexes involved in mRNA processing and export) that may prevent their diffusion back into the nucleolar-facing NE compartment. Our results establish *S. pombe* as a model for further studies determining the mechanisms that establish and maintain distinct populations of heterogeneous NPC composition within single nuclei. Importantly, our results demonstrate that the reduced NPC density and specific loss of Tpr-ortholog basket components over the nucleolus is not unique to budding yeast, but is a conserved feature of nuclear organization across highly divergent species. The ability for 3D-SIM to resolve and quantify individual NPCs labelled with multiple fluorescent proteins at endogenous levels provides tools to begin to interrogate how altered NPC compositions may allow for functional specialization of NPC function at distinct regions of the NE.

## Methods

### A. Yeast strains and plasmids

All *S. pombe* strains used in this manuscript are listed in **Table S2**. Deletion strains were obtained from the *S. pombe* haploid deletion library (Bioneer). Genes of interest were endogenously tagged using standard PCR-based methods (125), with lithium acetate transformation and colony selection as previously described (126). Cells were cultured in yeast extract with supplements (YES) media (5 g yeast extract, 30 g dextrose, 0.2 g each adenine, uracil, histidine, leucine and lysine, in 1 L of water) at 25°C, unless otherwise noted. For experiments using the *nmt41+* or *nmt3x* promoters, cells were cultured in Edinburgh minimal media with amino acid supplements (EMM5S) (127) at 30°C. Thiamine was added to EMM5S to a final concentration of 15 μM for 18-24 h at 30°C to repress expression. All strains were maintained in liquid culture for at least 48 h with back diluting to maintain cultures in logarithmic growth prior to imaging, unless otherwise noted. For growth assays, 4 OD_600_ of logarithmically growing cells were serially diluted ten-fold and spotted onto EMM5S agar plates (+/-15 μM thiamine) at 30°C. Where noted, cultures were treated with methyl benzimidazol-2-yl carbamate (MBC, 25 μg/mL), Cerulenin (10 μM) or dimethylsulfoxide (DMSO, vehicle control).

The coding sequence, or subdomains, for *lem2+* was amplified from genomic DNA using HiFi PCR master mix (Clontech) and cloned into NdeI/XhoI-digested pREP41-MCS+. The resulting plasmid was used as a template to amplify *nmt41-lem2-3xHA*, which was transformed into the *ura4+* locus of *lem2*Δ cells as described (128). Subdomains were similarly cloned and integrated. Integration was verified by PCR, and thiamine-dependent repression was validated by immuno blotting of whole cell extracts using anti-HA antibodies (Roche, 3F10).

### B. NPC quantitation and analysis by 3D-SIM

Exponentially growing cells were collected by centrifugation for 3 min at 3000 rcf and fixed in a solution of 4% formaldehyde supplemented with 200 mM glucose. Fixed cells were imaged in phosphate buffered saline, pH 7.4 with an Applied Precision OMX Blaze V4 (GE Healthcare) equipped with a 60x 1.42 NA Olympus Plan Apo oil objective and two PCO Edge sCMOS cameras. Two-color (GFP/mCherry) imaging was performed using 488-nm (GFP) or 561-nm (mCherry) lasers with alternating excitation, and a 405/488/561/640 dichroic with 504-552-nm and 590-628-nm emission filters. Images were acquired over a volume covering the entire nucleus with z-spacing of 125-nm (typically 4 μm). Widefield images of NPC clusters were acquired using the same microscope and settings as above but operating in widefield mode. SIM images were reconstructed with Softworx (Applied Precision Ltd), with a Wiener filter of 0.001. Except where noted, SIM images shown throughout are maximum intensity projections of all z-slices, scaled using bilinear interpolation with linear brightness and contrast adjustments in ImageJ (129).

Image analysis was performed using a number of custom plugins and macros for ImageJ/FIJI, all of which are freely available at http://research.stowers.org/imagejplugins/. Additional documentation and source code used for NPC density analysis can be found at http://www.stowers.org/research/ publications/libpb-1640. Statistical analysis was performed using R or Graph Pad Prism v 9.0. Average values along with standard deviation (SD) from the mean are shown based on the indicated number of samples (n), unless otherwise noted.

To quantitate the number of NPCs, individual nuclei were detected and segmented in an automated fashion using custom ImageJ plugins. Briefly, maximum intensity projections were used to perform automatic local thresholding for nuclear segmentation using a semi-automated protocol allowing the user to add and remove missed or poorly segmented ROIs. Each nucleus was cropped, and NPCs were detected using a “track max not mask” approach, in which the brightest voxel in the image is found and a spheroid with a diameter of 8 pixels (320 nm) in x and y and 5 slices in z (625 nm) is masked around that voxel. This process repeats until no voxels remain above a minimum threshold of 25% of the maximum intensity in the image. After NPC detection, the three-dimensional coordinates were used to model the NE surface using the “convhulln” function from the *geometry* package in R. Occasionally we observed the presence of points detected away from the NE (representing noise or foci of cytoplasmic signal). To remove these points prior to computation of the convex hull, we included an optimization step in which up to ten percent of the initial points could be removed if doing so increased the fraction of points present on the convex hull surface. Surface area and volume metrics were extracted for the 3D convex hull and used to derive NPC density values.

For a secondary method to validate density measurements, the nuclei image stacks were sum projected and background subtracted by selecting an ROI adjacent to the nucleus using the plugin “roi average subtract jru v1”. The resulting images were used to measure the nucleus area either by automated thresholding or occasionally by manual tracing of the nucleus in cases where thresholding failed to reliably segment the nucleus. The nucleus ROI areas and integrated densities were then extracted using “Analyze Particles” in ImageJ.

Cells were sorted into cell cycle stages using the following criteria: early G2, cell length < 9.5 μm, mononucleate; mid-G2, length between 9.5 and 11 μm, mononucleate; late G2/early mitosis, length >= 11.0 μm, mononucleate; late mitosis, length >= 11 μm, binucleate; G1/S, septated.

Single particle analysis was performed as previously described (85). Briefly, mother and daughter SPB spots were manually selected, and each spot was fitted to two 3D Gaussian functions and realigned along the axis between these two functions. To allow for visualization of NPC distributions in the x-y plane relative to the SPBs, a Euclidean distance filter was used in ImageJ to select for images in which both SPB points were at least 400 nm away from the edge of the nucleus based on maximum intensity projections. Realigned images were averaged as described previously (84, 85). To account for biologically irrelevant directional bias in the x-y plane, the averaged images were further averaged with a mirrored (x-y) image. All averaged images presented were thresholded to display pixels above a threshold of 25% of the maximum intensity value. Quantitation of Nup intensity relative to the mother and daughter SPB was performed in an automated manner, using line profiles with a width (12 pixels) covering both mother and daughter SPBs to generate profiles of the Nup/GFP intensities. Proteins of interest were considered to be excluded if the normalized peak intensity fell below 0.8, based on the reduction in signal observed in simulated random datasets. For excluded proteins, the plot profiles were fit to a Gaussian and the width of the exclusion zone was measured by computing the full-width half-maximal (FWHM) value for the fit curve using the equation FWHM = 2.355 x standard deviation (SD). The size of exclusion zones for proteins of interest were compared using the FWHM value from plot profiles of the averaged SPA-SIM images, with error bars representing the 95% confidence interval of the SD of the Gaussian fit. Curve fitting and statistical analyses were performed using GraphPad Prism v. 9.0.

An analogous approach was used to generate an averaged image for Nups relative to the SPB in anaphase and telophase nuclei. First, nuclei in these stages were identified in which the SPB was clearly observed within the NE from single z-slices. These slices were then used to manually trace the NE (based on Nup signal), followed by straightening of the polyline using the ImageJ macro “polyline profile jru v1” with a line width of 8 pixels and the option for “Output Straightened” selected. The resulting straightened images were used to identify the location of the SPB, and a 1 μm region of the image centered on the SPB was cropped. The cropped images were then combined, and an averaged image was generated. The averaged image was further averaged with a mirrored image (in both horizontal and vertical directions) to generate a final averaged image with no directional biases.

### C. Simulations and modeling

For comparison of SPA-SIM data to the distribution expected to be observed by random chance, we simulated spherical nuclei with a of radius 1.25 μm (based on average dimensions of mid-G2 stage nuclei) with 125 randomly positioned NPCs to model an NPC density (6.4 NPCs/μm^2^) similar to the average NPC density observed by 3D-SIM. NPCs were simulated as 3D Gaussians with a FWHM in the x-y plane of 100 nm and 300 nm in z, a minimum center-to-center distance of 100 nm and a maximum intensity of 100 photons. The simulated pixel size was 40 nm with a z-slice spacing of 125 nm. The resulting intensities were multiplied by 20 (artificial “Gain”) and a Gaussian read noise with a standard deviation of 40 intensity units was added to each voxel. For simulation of SPA-SIM data, two SPB points were simulated with a center-to-center distance of 180 nm in a second channel. Simulated images were processed using the same approaches outlined above for NPC quantitation and SPA-SIM analyses.

### D. Confocal imaging

Confocal imaging to determine NPC density over the nucleolus was performed in log-phase cells expressing the nucleolar protein, Nuc1-mCherry, and utilized a PerkinElmer UltraVIEW VoX with a Yokogawa CSU-X1 spinning disk head, a 100x 1.46 NA Olympus Plan Apo oil objective and CCD (ORCA-R2) and EMCCD (C9100-13) cameras. GFP/mCherry images were taken using a 488-nm laser (for GFP/mNeonGreen) or a 561-nm laser (for mCherry), with alternating excitation. Images were collected using the Volocity imaging software with a z spacing of 0.3 μm over a volume of 8 μm. To assess nucleoporin intensity levels at the nuclear envelope, the middle four slices of the image stack were sum projected using ImageJ, and the nuclear envelope was manually traced to generate line profiles for both GFP (Nup) and mCherry (nucleolus) channels in nuclei where the nucleolus was oriented to one side of the nucleus in the x-y direction. The line profiles for Nuc1-mCherry were boxcar smoothened, thresholded at their half-maximal values, and all profiles were aligned at the center of the Nuc1-mCherry peak. The nucleoporin line profiles were then resampled and averaged to generate average intensity profiles. Mean normalized intensity values for the Nup signal at the center of the nucleolar peak were calculated from three independent biological replicates and plotted in GraphPad Prism v. 9.0.

To analyze spindle morphology and NPC cluster dynamics in live cells, *≈* 200-300 μL of log phase cells were applied to 35-mm glass bottom dishes (MaTek, no. 1.5 coverslip) that had been pre-coated with 1 mg/mL soybean lectin (in water) for 15 min and rinsed with YES media. After cells were allowed to settle for 30 min at 30°C, 2 mL of pre-warmed YES media was carefully added. Cells were imaged on a Nikon Ti-E microscope equipped with a CSI W1 spinning disk (Yokogawa) using a 60x 1.4 NA Olympus Plan Apo oil objective and an iXon DU897 Ultra EMCCD (Andor) camera. GFP and mCherry were excited at 488 nm and 561 nm, respectively, and collected through ET525/36m (GFP) or ET605/70m (mCherry) bandpass filters. Samples were maintained at 30°C using an Oko Lab stage top incubator. Images were acquired overa6 μm volume with 0.3 μm z-spacing for 45 min at 2 min intervals.

SPB distances were measured using the Euclidean distance between manually annotated SPBs from maximum intensity projections of the full image stacks. Spindle orientations were determined manually using the Angle tool in ImageJ to measure the difference of the spindle relative to the cell axis from maximum intensity projected image stacks. To generate image montages presented in figures, maximum intensity projections of the full image stacks were background subtracted, bleach corrected (using the Simple Ratio method) and scaled 3-fold (x and y) using bilinear interpolation.

### E. Electron microscopy

The distance between SPBs and the nearest NPC was measured in images from samples prepared as previously described (85). Sections in which both the outer and inner nuclear membranes were clearly resolved were used for analysis. The distance from the center of the SPB to the nearest NPC (determined based on visible fusion of the INM and ONM) was measured by manually tracing the nuclear envelope using the polyline tool in ImageJ, and the data was plotted in GraphPad Prism v. 9.0.

## ACKNOWLEDGEMENTS

We thank Julie Cooper for strains, Zulin Yu and members of the Stowers Microscopy Core Facility for imaging assistance, and Brian Slaughter and members of the Jaspersen lab for discussion and feedback on this manuscript. Schematics throughout were created using BioRender.com. Research reported in this publication was supported by the Stowers Institute for Medical Research and NIH-NIGMS under award number R01GM121443 (to SLJ). J.M.V is a recipient of a Ruth L. Kirschstein NRSA Postdoctoral Fellowship (F32GM133096). Original data underlying this manuscript can be downloaded from the Stowers Original Data Repository at http://www.stowers.org/research/publications/libpb-1640. The authors declare no competing financial interests.

## DATA AVAILABILITY STATEMENT

Upon peer-reviewed publication, all strains generated for and used in this study will be made available upon reasonable request. Original data underlying this manuscript will be accessible at the Stowers Original Data Repository at http://www.stowers.org/research/publications/libpb-1640.

---

**Fig. S1.**
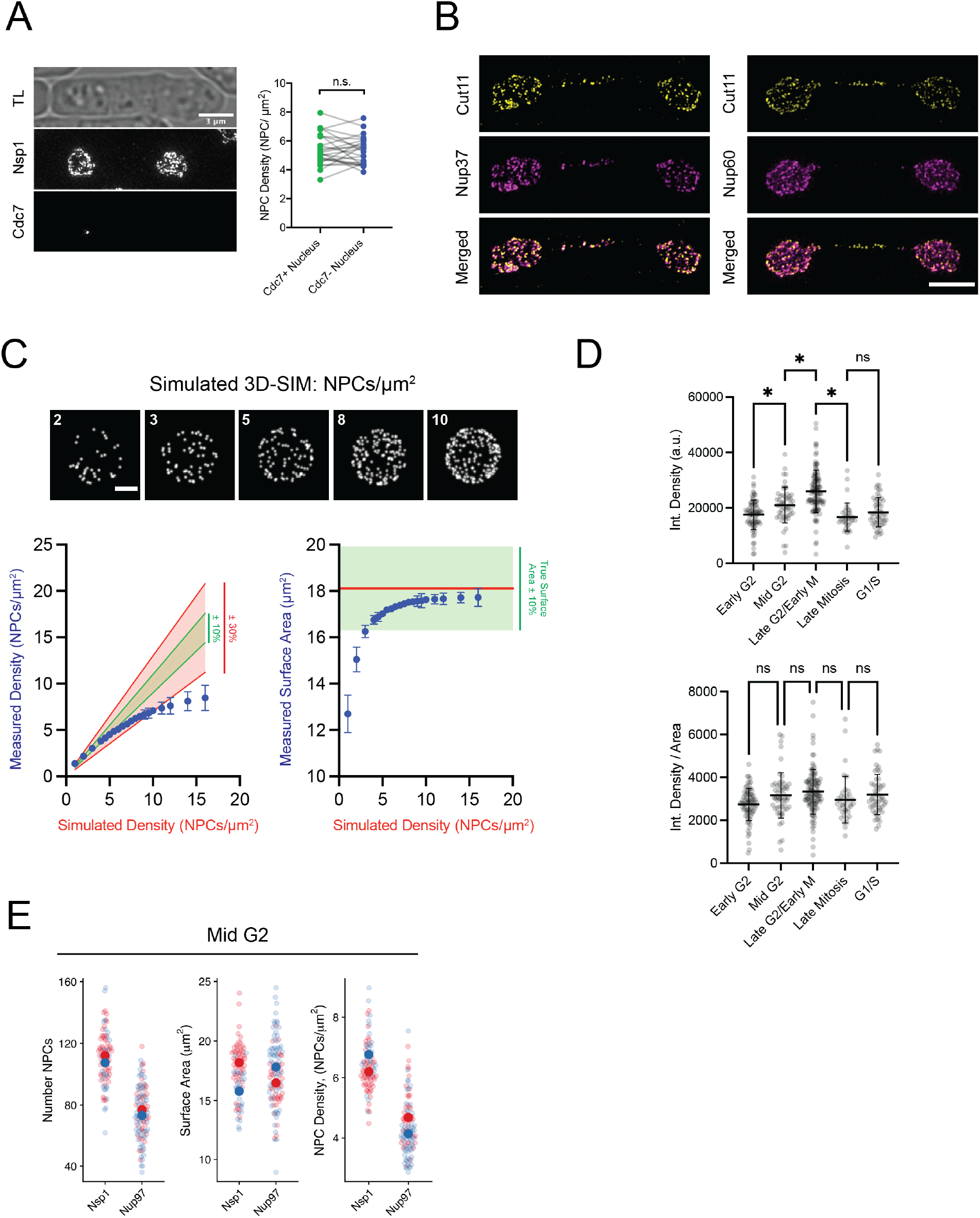
NPC analysis. A) 3D-SIM of Nsp1-mCh NPCs and Cdc7-GFP, which marks the “new” SPB. Quantitation of NPC density in nuclei that did and did not inherit Cdc7-GFP. Bar, 3 μm. ns, non-significant differences determined by Wilcoxon matched-pairs signed rank test. B) 3D-SIM projections of late anaphase nuclei with Cut11-GFP (yellow) and Nup37-mCh or Nup60-mCh (magenta). Bar, 3 μm. Loss of Nup60-GFP in the midzone region is due to NPC disassembly (Dey et al, 2020; Expósito-Serrano, et al., 2020). C) Projections of simulated 3D NPC distributions at densities ranging from 2-10 NPCs/μm^2^. Bar, 1 μm. Graphs comparing the true simulated NPC density and surface areas with the values obtained from the NPC analysis pipeline outlined in Figure 1C. Predicted ranges for values with 10% (green) and/or 30% (red) error ranges are highlighted. D) Quantitation of total Nsp1-GFP signal (integrated density) (top), and signal normalized to the nuclear area, validating increases in Nsp1-GFP NPCs through the cell cycle while NPC density is maintained. Asterisk indicates significant difference in mean values determined by One-Way ANOVA with Šidák’s multiple comparisons test. E) Comparison of NPC number and densities using Nsp1-GFP and Nup97-GFP.

**Fig. S2.**
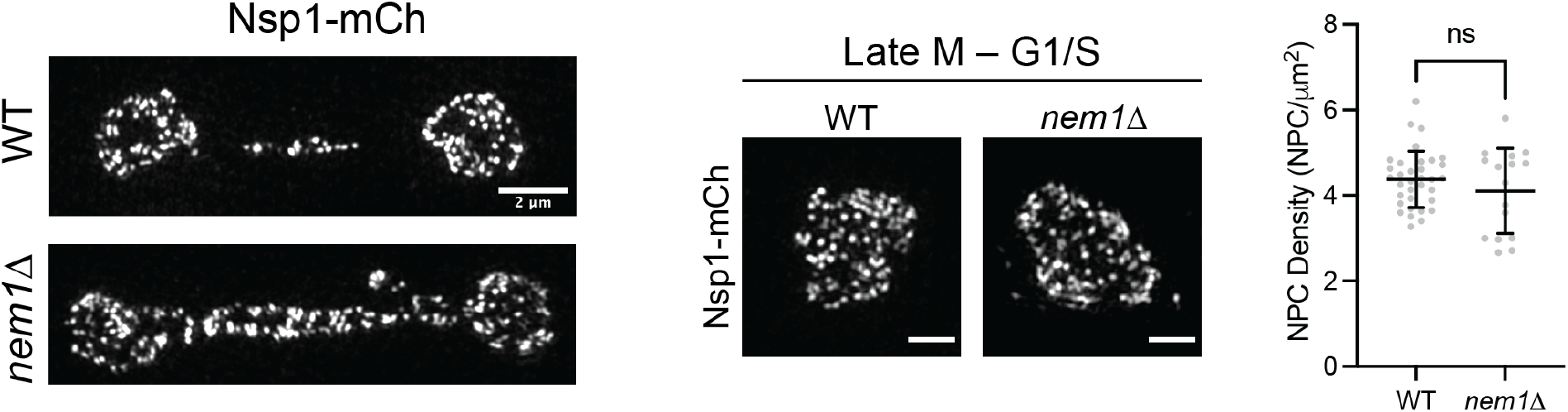
NPC sequestration to the anaphase bridge in *nem1*Δ. 3D-SIM of Nsp1-mCh NPCs in wild-type and *nem1*Δ nuclei in late anaphase. Bar, 2 μm. *nem1*Δ nuclei have normal morphologies and equivalent NPC densities as wild-type nuclei after completing mitosis (Late M and G1/S, determined by Mann-Whitney test). Bar, 1 μm.

**Fig. S3.**
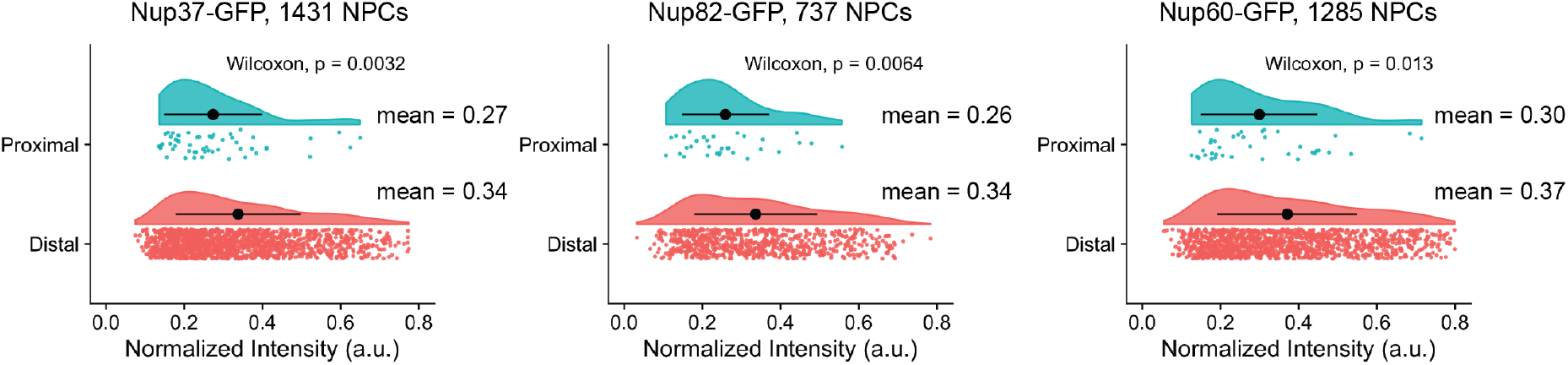
Nup intensity proximal to the SPB. Nup-GFP intensity in SPB-proximal (<100 nm) and distal (>100 nm) regions. Values in each were compared using a Wilcoxon rank sum test. Black dots represent the mean normalized intensity value, and error bars show SD.

**Fig. S4.**
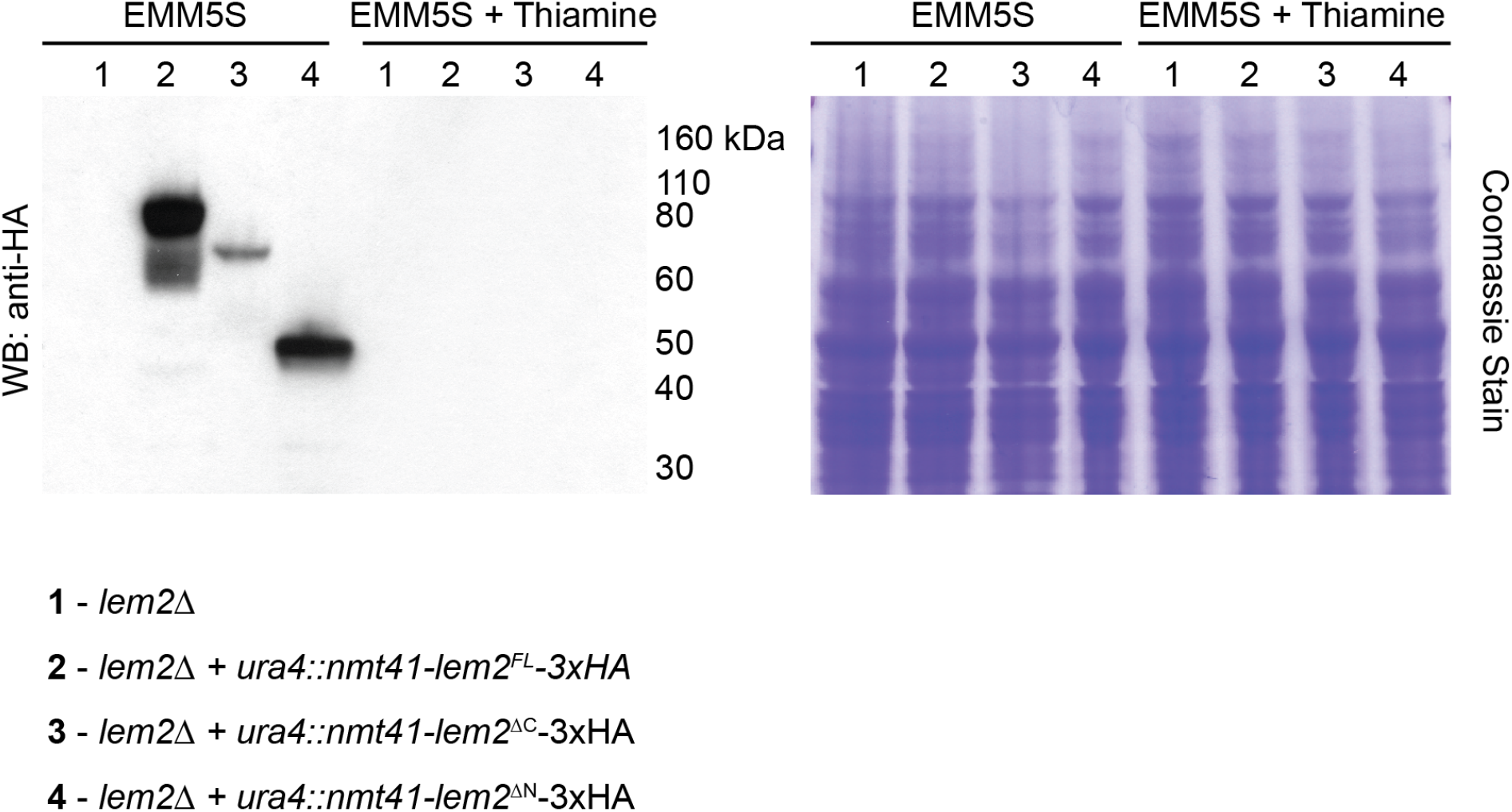
Expression of Lem2 rescue constructs. Immunoblot of strains from Figure 6D. Expression of each construct was observed for each fusion protein by anti-HA immuno blotting in cells cultured in EMM5S (no thiamine) and is reduced to undetectable levels after 22 hours of culture in EMM5S with 15 μM thiamine. A Coomassie stained gel is shown as a loading control. The position of molecular weight markers (kDa) is shown.

## Supplemental Tables

Summary statistics from four independent biological replicates of Nsp1-GFP 3D-SIM imaging experiments. Stages assigned as described in Materials and Methods. Surface Area, Volume, Sphericity and NPC Density values derived from the computed 3D convex hull. Points Removed represents the average number of NPC points removed during complex hull optimization as described in Materials and Methods.

**Table S1:** Summary statistics for Nsp1-GFP NPC analysis.

| Stage | Cell Length (μm) | Surface Area (μm<sup>2</sup>) | Volume (μm<sup>3</sup>) | Sphericity | Number NPCs | NPC Density (NPCs per μm<sup>2</sup>) | Points Removed | N |
| --- | --- | --- | --- | --- | --- | --- | --- | --- |
| Early G2 | 8.7 ± 0.5 | 16.9 ± 2.7 | 5.9 ± 1.4 | 0.93 ± 0.02 | 95.3 ± 19.9 | 5.7 ± 1.0 | 0.8 ± 1.2 | 233 |
| Mid G2 | 10.2 ± 0.4 | 18.4 ± 2.8 | 6.7 ± 1.6 | 0.93 ± 0.02 | 105.1 ± 20.2 | 5.8 ± 1.0 | 0.9 ± 1.5 | 174 |
| Late G2/<br>Early M | 12.7 ± 1.2 | 21.8 ± 4.1 | 8.7 ± 2.4 | 0.93 ± 0.02 | 123.7 ± 26.6 | 5.8 ± 1.2 | 0.9 ± 1.3 | 317 |
| Late M | 14.0 ± 0.9 | 14.6 ± 2.6 | 4.6 ± 1.3 | 0.91 ± 0.02 | 77.5 ± 13.9 | 5.4 ± 1.1 | 1.2 ± 1.8 | 122 |
| G1/S | 14.2 ± 1.0 | 15.5 ± 2.8 | 5.1 ± 0.5 | 0.92 ± 0.02 | 81.1 ± 17.1 | 5.3 ± 0.9 | 0.9 ± 1.2 | 162 |

**Table S2:** Yeast strains.

| Strain | Genotype | Figure | Source/Reference |
| --- | --- | --- | --- |
| fySLJ707 | *h-, nsp1-GFP-KanMX6, his3-D1, leu1-32, ura4-D18, ade6-M210* | Fig. 1A-E, S1D, S1E | This study |
| fySLJ485 | *h-, nup146-GFP-KanMX6, his3-D1, leu1-32, ura4-D18, ade6-M210* | Fig. 1B | This study |
| fySLJ748 | *h+, nup37-GFP-KanMX6, his3-D1, leu1-32, ura4-D18, ade6-M210* | Fig. 1B | This study |
| fySLJ484 | *h-, nup97-GFP-KanMX6, his3-D1, leu1-32, ura4-D18, ade6-M210* | Fig. 1B, S1E | This study |
| fySLJ035 | *h-, cut11::GFP:ura4+, leu1-32, ura4-D18* | Fig. 1B | J.R. McIntosh, (Mcl316) |
| fySLJ730 | *h-, nup60-GFP-KanMX6, his3-D1, leu1-32, ura4-D18, ade6-M210* | Fig. 1B | This study |
| fySLJ714 | *h-, cdc7-GFP-KanMX6, nsp1-mCherry-NatMX6, his3-D1, leu1-32, ura4-D18, ade6-M210* | Fig. S1A | This study |
| fySLJ771 | *h?, cut11::GFP:ura4+, nup37-mCherry-NatMX6, leu1-32, ura4-D18* | Fig. S1B | This study |
| fySLJ772 | *h?, cut11::GFP:ura4+, nup60-mCherry-NatMX6, leu1-32, ura4-D18* | Fig. S1B | This study |
| fySLJ457 | *h+, nsp1-mCherry-NatMX6, his3-D1, leu1-32, ura4-D18, ade6-M210* | Fig. 2A, 3 | This study |
| fySLJ944 | *h+, wee1-50, nup37-mCherry-HygMX6, ade6-M210, leu1-32, ura4-D18* | Fig. 2B | This study |
| fySLJ1118 | *h?, ppc89-mCherry-NatMX6, nsp1-GFP-KanMX6, cdc25.22, leu1-32, ura4-D18* | Fig. 2C | This study |
| fySLJ541 | *h-, nsp1-mCherry-NatMX6, (prototroph)* | Fig. 2D | This study |
| fySLJ1135 | *h?, atg1Δ::KanMX6, nsp1-mCherry-NatMX6 (prototroph)* | Fig. 2D | This study |
| fySLJ1136 | *h?, atg8Δ::KanMX6, nsp1-mCherry-NatMX6 (prototroph)* | Fig. 2D | This study |
| fySLJ515 | *h?, nup132Δ::KanMX6, nsp1-mCherry-NatMX6, leu1-32, ura4-D18* | Fig. 3A-C | This study |
| fySLJ879 | *h?, nem1Δ::KanMX6, nsp1-mCherry-NatMX6, leu1-32, ura4-D18* | Fig. 3D-E, S2 | This study |
| fySLJ1147 | *h?, nup132Δ::KanMX6, nsp1-mCherry-NatMX6, Sad1-GFP-NatMX6* | Fig. 3F,G | This study |
| fySLJ712 | *h?, nuc1-mCherry-NatMX6, nup44-GFP-KanMX6, his3-D1, leu1-32, ura4-D18, ade6-M210* | Fig. 4A-D | This study |
| fySLJ713 | *h?, nuc1-mCherry-NatMX6, nup146-GFP-KanMX6, his3-D1, leu1-32, ura4-D18, ade6-M210* | Fig. 4B-D | This study |
| fySLJ997 | *h?, nuc1-mCherry-NatMX6, nup124-GFP-KanMX6, his3-D1, leu1-32, ura4-D18, ade6-M210* | Fig. 4B-D | This study |
| fySLJ998 | *h-, nuc1-mCherry-NatMX6, nup211-GFP-KanMX6, his3-D1, leu1-32, ura4-D18, ade6-M210* | Fig. 4B-D | This study |
| fySLJ1001 | *h-, nuc1-mCherry-NatMX6, nup97-mNeonGreen-HygMX6, his3-D1, leu1-32, ura4-D18, ade6-M210* | Fig. 4B-D | This study |
| fySLJ1017 | *h?, nuc1-mCherry-NatMX6, nup60-GFP-KanMX6, his3-D1, leu1-32, ura4-D18, ade6-M210* | Fig. 4B-D | This study |
| fySLJ1020 | *h?*, *nuc1-mCherry-NatMX6*, *nup61-GFP-KanMX6*, *his3-D1*, *leu1-32*, *ura4-D18*, *ade6-M210* | Fig. 4B-D | This study |
| fySLJ1021 | *h?*, *nuc1-mCherry-NatMX6*, *alm1-GFP-KanMX6*, *his3-D1*, *leu1-32*, *ura4-D18*, *ade6-M210* | Fig. 4B-D | This study |
| fySLJ1022 | *h?*, *nuc1-mCherry-NatMX6*, *seh1-GFP-KanMX6*, *his3-D1*, *leu1-32*, *ura4-D18*, *ade6-M210* | Fig. 4B-D | This study |
| fySLJ1208 | *h+*, *les1-mNeonGreen-HygMX6*, *ppc89-mCherry-NatMX6*, *his3-D1*, *leu1-32*, *ura4-D18*, *ade6-M210* | Fig. 5B | This study |
| fySLJ738 | *h?*, *nsp1-GFP-KanMX6*, *ppc89-mCherry-NatMX6*, *his3-D1*, *leu1-32*, *ura4-D18*, *ade6-M210* | Fig. 5B-G, 7B-C | This study |
| fySLJ1122 | *h+*, *sad1-GFP-KanMX6*, *adh15::mCherry-atb2-NatMX6* | Fig. 5C | This study |
| fySLJ745 | *h-*, *nup60-GFP-KanMX6*, *ppc89-mCherry-NatMX6*, *his3-D1*, *leu1-32*, *ura4-D18*, *ade6-M210* | Fig. 5F-G, S3 | This study |
| fySLJ746 | *h+*, *nup82-GFP-KanMX6*, *ppc89-mCherry-NatMX6*, *his3-D1*, *leu1-32*, *ura4-D18*, *ade6-M210* | Fig. 5F-G, S3 | This study |
| fySLJ747 | *h+*, *nup85-GFP-KanMX6*, *ppc89-mCherry-NatMX6*, *his3-D1*, *leu1-32*, *ura4-D18*, *ade6-M210* | Fig. 5F-G | This study |
| fySLJ822 | *h90*, *nup37-GFP-KanMX6*, *ppc89-mCherry-NatMX6*, *his3-D1*, *leu1-32*, *ura4-D18*, *ade6-M210* | Fig. 5F-G, S3 | This study |
| fySLJ530 | *h?*, *nup97-GFP-KanMX6*, *ppc89-mCherry-NatMX6*, *his3-D1*, *leu1-32*, *ura4-D18*, *ade6-M210* | Fig. 5F-G | This study |
| fySLJ867 | *h?*, *alm1-GFP-KanMX6*, *ppc89-mCherry-NatMX6*, *his3-D1*, *leu1-32*, *ura4-D18*, *ade6-M210* | Fig. 5F-G | This study |
| fySLJ577 | *h+*, *lem2-GFP-KanMX6*, *ppc89-mCherry-NatMX6*, *his3-D1*, *leu1-32*, *ura4-D18*, *ade6-M210* | Fig. 6A | This study |
| fySLJ845 | *h?*, *nsp1-GFP-KanMX6*, *ppc89-mCherry-NatMX6*, *ima1Δ::HygMX6*, *leu1-32*, *ura4-D18* | Fig. 6B | This study |
| fySLJ849 | *h?*, *nsp1-GFP-KanMX6*, *ppc89-mCherry-NatMX6*, *lem2Δ::HygMX6*, *leu1-32*, *ura4-D18* | Fig. 6B | This study |
| fySLJ860 | *h?*, *nsp1-GFP-KanMX6*, *ppc89-mCherry-NatMX6*, *man1Δ::HygMX6*, *leu1-32*, *ura4-D18* | Fig. 6B | This study |
| fySLJ1209 | *h?*, *nsp1-GFP-KanMX6*, *ppc89-mCherry-NatMX6*, *csi1Δ::HygMX6*, *leu1-32*, *ura4-D18* | Fig. 6E | This study |
| fySLJ1042 | *h+*, *bleMX6-nmt3X-bqt1-GBP-mCherry:HygMX6* | Fig. 7C | This study |
| fySLJ1104 | *h?*, *bleMX6-nmt3X-bqt1-GBP-mCherry:HygMX6*, *nsp1-GFP-KanMX6*, *ppc89-mCherry-NatMX6* | Fig. 7B,C | This study |
| fySLJ1124 | *h-*, *bleMX6-nmt3X-bqt1-GBP-mCherry:HygMX6*, *nsp1-GFP-KanMX6*, *ppc89-mCherry-NatMX6*, *adh15::mCherry-atb2-NatMX6* | Fig. 7D-E | This study |
| fySLJ1117 | *h?*, *nsp1-GFP-KanMX6*, *ppc89-mCherry-NatMX6*, *lem2Δ::HygMX6*, *ura4::nmt41-lem2FL-3xHA-ura4+*, *leu1-32*, *ura4-D18* | Fig. 6D, S4 | This study |
| fySLJ1142 | *h?*, *nsp1-GFP-KanMX6*, *ppc89-mCherry-NatMX6*, *lem2Δ::HygMX6*, *ura4::nmt41-lem2ΔC-3xHA-ura4+*, *leu1-32*, *ura4-D18* | Fig. 6D, S4 | This study |
| fySLJ1144 | *h?*, *nsp1-GFP-KanMX6*, *ppc89-mCherry-NatMX6*, *lem2Δ::HygMX6*, *ura4::nmt41-lem2ΔN-3xHA-ura4+*, *leu1-32*, *ura4-D18* | Fig. 6D, S4 | This study |

